# Mutation-induced biophysical destabilization as a key contributor to cancer-driving potential in the human structural protein interactome

**DOI:** 10.64898/2026.04.15.718717

**Authors:** Ting-Yi Su, Yu Xia

**Affiliations:** Graduate Program in Quantitative Life Sciences, McGill University, Montréal, Québec, Canada; Department of Bioengineering, McGill University, Montréal, Québec, Canada

**Keywords:** human structural protein interactome, cancer, cancer-driving potential, protein-folding stability, protein-binding stability, destabilizing mutations, loss of function, structural systems biology

## Abstract

A systems-level investigation of mutation-induced perturbations in the human structural protein interactome, the network of structurally resolved protein–protein interactions in human cells, provides mechanistic insight into the complexity underlying oncogenesis. Mutations can destabilize protein folding or specific protein–protein interactions, resulting in loss-of-function effects within the interactome. Although such network-level loss-of-function consequences may contribute to oncogenesis, previous studies have largely examined either folding or binding destabilization in isolation. Here, we performed structural and free-energy calculations to assess the impact of interactome-wide cancer-associated missense mutations on protein-folding stability and protein-binding stability. We assessed the cancer-driving potential of destabilizing mutations using a “fold difference” metric, defined as the ratio between the fractions of destabilizing mutations in datasets of cancer-associated and non-pathogenic mutations. We observed a strong positive correlation between biophysical destabilization and cancer-driving potential, with folding-destabilizing (“quasi-null”) and binding-destabilizing (“edgetic”) mutations within cancer-driving genes causing stronger structural and functional effects than those across the entire cancer genome. Our findings align with the expectation that cancer-driving genes are enriched in driver mutations and suggest that biophysical destabilization is a key contributor to cancer-driving potential. Overall, our study provides a biophysical perspective on the loss-of-function implications of destabilizing mutations across the interactome, enhancing comprehension of the intricate processes driving oncogenesis.

## Introduction

Cancer, a highly heterogeneous disease resulting from an accumulation of somatic mutations across multiple genes, can be better understood by studying mutations that confer cancer-driving potential. In particular, a systems-level approach involving the human protein interactome, the network of all protein-protein interactions (PPIs) in human cells, has been introduced for studying genotype-to-phenotype associations in cancer (Yi et al., 2017). Several large-scale efforts have mapped cancer-associated mutations onto the human structural protein interactome, focusing on mutations that overlap with important structural elements or PPI interfaces (J. Zhang, Pei, Durham, Bos, & Cong, 2022; Y. Zhang et al., 2025). Such mutations are more likely to result in functional consequences contributing to cancer-driving potential. Thus, investigating the biophysical implications of these mutations across the human structural protein interactome provides mechanistic insight into genotype-to-phenotype relationships and the processes driving oncogenesis.

Although cancer is often described as a proliferative disease, mutations can disrupt and reshape the network of PPIs that underlie cellular function, resulting in loss-of-function (LoF) effects within the interactome (Fu, Mo, & Ivanov, 2025; Sharifi Tabar, Francis, Yeo, Bailey, & Rasko, 2022; Shi & Moult, 2011). Such network-level LoF effects manifest as the loss of nodes, when entire proteins are destabilized or degraded, or as the removal of edges, when specific PPIs are destabilized. While studies have employed computational approaches to investigate protein-folding (folding ΔΔG) (Chillón-Pino, Badonyi, Semple, & Marsh, 2024; Shi & Moult, 2011) or protein-binding (binding ΔΔG) (Engin, Kreisberg, & Carter, 2016; Nishi et al., 2013) destabilization of cancer-associated mutations, these LoF effects have generally been investigated in isolation. To the best of our knowledge, no study has yet computationally calculated and investigated both folding ΔΔG and binding ΔΔG. As a result, despite the growing structural coverage of the human protein interactome, the interactome-wide LoF consequences of destabilizing mutations in cancer have been largely unexplored.

While destabilizing mutations can drive oncogenesis, determining which of these mutations are true drivers remains challenging. Distinguishing driver mutations from passenger mutations, neutral changes that do not contribute to oncogenesis, is complicated by the vast number of somatic mutations and the genetic heterogeneity both within tumors and across patients (Kumar et al., 2020; Ostroverkhova, Przytycka, & Panchenko, 2023). Furthermore, not all mutations in a gene classified as cancer-driving are functionally relevant; some may have unknown significance and could in fact be passenger mutations (Porta-Pardo, Valencia, & Godzik, 2020). Studies have shown that specific cancer-associated mutations disrupting protein folding or PPIs remodel interactome topology, driving changes in cancer-related biochemical processes (Mehnert et al., 2020; Xiong, Lee, Li, Zhao, & Yu, 2022). In this context, calculating the folding ΔΔG and binding ΔΔG of interactome-wide mutations enables a systems-level quantification of the functional impact of destabilizing mutations. This approach highlights a subset of functionally relevant somatic mutations and offers a biophysical perspective for assessing cancer-driving potential.

The functional and mechanistic insights obtained from studying destabilizing mutations in Mendelian disorders provide a valuable framework for interpreting similar mutations in cancer (Haque et al., 2025; Kaminker et al., 2007; Melamed, Emmett, Madubata, Rzhetsky, & Rabadan, 2015; Song et al., 2023). Mendelian diseases arise from germline mutations in single genes and are often associated with highly penetrant phenotypes. In contrast, cancer is driven by somatic mutations across multiple genes, thus the effects of individual mutations are usually partial and influenced by genetic, cellular, and epigenetic context (Rahman, 2014; Taeubner et al., 2018). Nonetheless, in both disease types, destabilizing mutations can manifest as LoF effects within the interactome, suggesting that insights from Mendelian diseases can guide interpretation of the more heterogeneous and context-dependent implications in cancer. We recently demonstrated that, in Mendelian diseases, mutation-induced protein-folding destabilization correlates with a “fold difference” metric of functional and phenotypic deleteriousness (Su & Xia, 2025). This finding suggests that our fold difference metric, built upon observations from Mendelian diseases, can be applied to evaluate the cancer-driving potential of destabilizing mutations.

In this work, we structurally resolved the human protein interactome at the atomic level and performed a ΔΔG-based assessment of interactome-wide mutations to investigate the potential for destabilizing mutations to drive oncogenesis. We obtained cancer-associated mutations across the entire cancer genome and within cancer-driving genes. We mapped mutations onto the interactome and performed structural and free-energy calculations to evaluate their biophysical implications. We calculated the effects of missense mutations on protein-folding stability, defined as folding ΔΔG (folding ΔG_mutant_ – folding ΔG_wildtype_), and on protein-binding stability, defined as binding ΔΔG (binding ΔG_mutant_ – binding ΔG_wildtype_). We focused on the network-level LoF effects of folding-destabilizing (“quasi-null”) and binding-destabilizing (“edgetic”) mutations (**Figure 1**). We adopted our “fold difference” metric (Su & Xia, 2025) for quantifying the functional and phenotypic deleteriousness of Mendelian disease-causing quasi-null mutations to study the cancer-driving potential of quasi-null and edgetic mutations. We defined the fold difference for quasi-null mutations as the ratio between the fractions of quasi-null mutations in datasets of cancer-associated and non-pathogenic missense mutations. Similarly, we defined the fold difference for edgetic mutations as the ratio between the fractions of edgetic mutations in datasets of cancer-associated and non-pathogenic missense mutations. We observed that protein-folding destabilization correlates with the deleteriousness of cancer-associated quasi-null mutations, with quasi-null mutations within cancer-driving genes exhibiting stronger structural and functional effects than those across the entire cancer genome. Similarly, we observed that protein-binding destabilization correlates with the deleteriousness of cancer-associated edgetic mutations, with edgetic mutations within cancer-driving genes exhibiting stronger structural and functional effects than those across the entire cancer genome. Altogether, these observations suggest that biophysical destabilization, of both folding and binding, is a key contributor to cancer-driving potential. Overall, our study of interactome-wide destabilizing mutations provides a comprehensive biophysical understanding of the complex mechanisms underlying oncogenesis.

**Figure 1.**
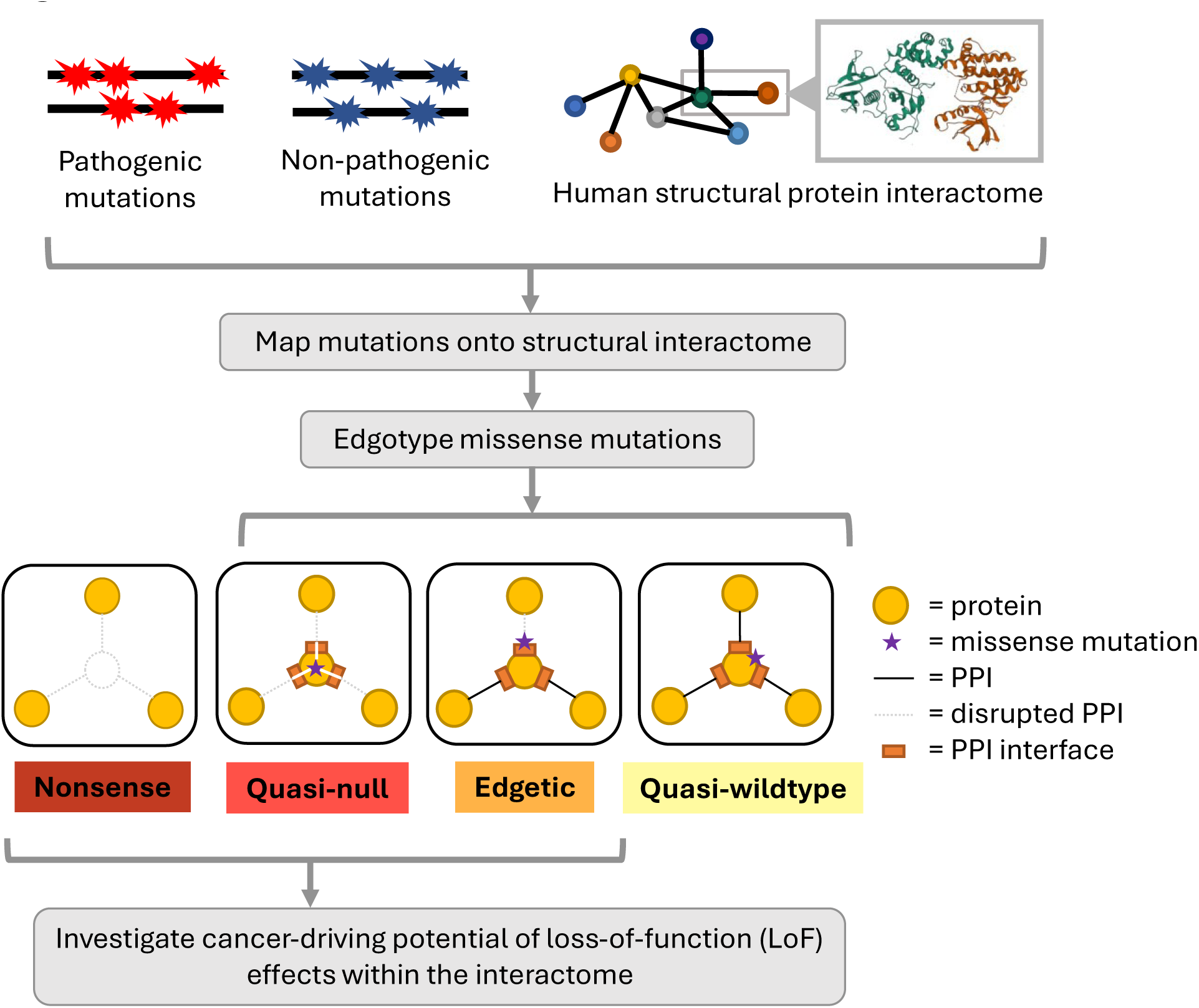
Pipeline for investigating the cancer-driving potential of missense (quasi-null and edgetic) and nonsense mutations associated with loss-of-function (LoF) effects within the interactome. Quasi-null (folding-destabilizing) and nonsense mutations result in node removals, while edgetic (binding-destabilizing) mutations result in edge removals.

## Results

### Constructing human structural protein interactomes and mapping mutations

We constructed two human structural interactomes (SIs): HI-union-SI and IntAct-SI. The HI-union-SI was built using the HI-union database of experimentally-determined PPIs (Luck et al., 2020), while the IntAct-SI was built using the IntAct database of literature-curated PPIs (del Toro et al., 2022). Protein sequences and chain sequences in three-dimensional (3D) protein structures were obtained from the UniProtKB/Swiss-Prot (The UniProt Consortium, 2023) and the Protein Data Bank (PDB) (Berman et al., 2000) databases, respectively. We constructed 3D homology models of PPIs within our two SIs using MODELLER (Webb & Sali, 2016) and BLASTP (Camacho et al., 2009) alignments of Swiss-Prot proteins against homologous PDB chains. Our HI-union-SI comprises 2,195 structural PPIs among 1,704 Swiss-Prot proteins, while our IntAct-SI comprises 5,490 structural PPIs among 4,004 Swiss-Prot proteins.

Within nonsynonymous mutations, we specifically investigated missense and nonsense mutations. Non-pathogenic missense and nonsense mutations were curated from dbSNP (Sherry et al., 2001). Cancer-associated missense and nonsense mutations were curated from whole-genome screens of cancer patients (COSMIC genome screen) (Sondka et al., 2024) and from genes in the Cancer Gene Census (COSMIC CGC), a literature-curated and experimentally-validated dataset of cancer-driving genes (Sondka et al., 2018). We investigated both the COSMIC genome screen and the COSMIC CGC to compare the cancer-driving potential of mutations in each dataset. Mendelian disease-causing missense and nonsense mutations were curated from ClinVar (Landrum et al., 2016). Mutations were mapped onto our SIs using transcript to protein mappings from RefSeq (O’Leary et al., 2016) and Ensembl (Martin et al., 2023), and were retained only if they were covered by at least one PPI structural model. The HI-union-SI contains 8,187 dbSNP, 44,680 COSMIC genome screen, 4,385 COSMIC CGC, and 462 ClinVar missense mutations, as well as 164 dbSNP, 3,018 COSMIC genome screen, 336 COSMIC CGC, and 348 ClinVar nonsense mutations. The IntAct-SI contains 23,128 dbSNP, 128,212 COSMIC genome screen, 18,486 COSMIC CGC, and 1,449 ClinVar missense mutations, as well as 378 dbSNP, 8,356 COSMIC genome screen, 1,655 COSMIC CGC, and 1,688 ClinVar nonsense mutations.

### Predicting binding-destabilizing and folding-destabilizing missense mutations

We predicted each missense mutation to be either quasi-wildtype (non-destabilizing), edgetic (binding-destabilizing), or quasi-null (folding-destabilizing) based on its location within the protein structure and its FoldX 5.1 (Delgado et al., 2025; Schymkowitz et al., 2005) binding ΔΔG or folding ΔΔG. We chose FoldX over other mutagenesis tools for its computational efficiency and ease of use. An edgetic mutation is located at an interfacial residue and predicted to disrupt the binding stability of at least one associated PPI (binding ΔΔG ≥0.8 kcal/mol). A quasi-null mutation is predicted to disrupt the folding stability of the protein (folding ΔΔG ≥2 kcal/mol). The mean BLOSUM62 scores (Henikoff & Henikoff, 1992) of the amino acid substitutions decreased progressively from quasi-wildtype to edgetic to quasi-null mutations (**Figure 2**, bottom row), supporting our predictions of their functional impact.

**Figure 2.**
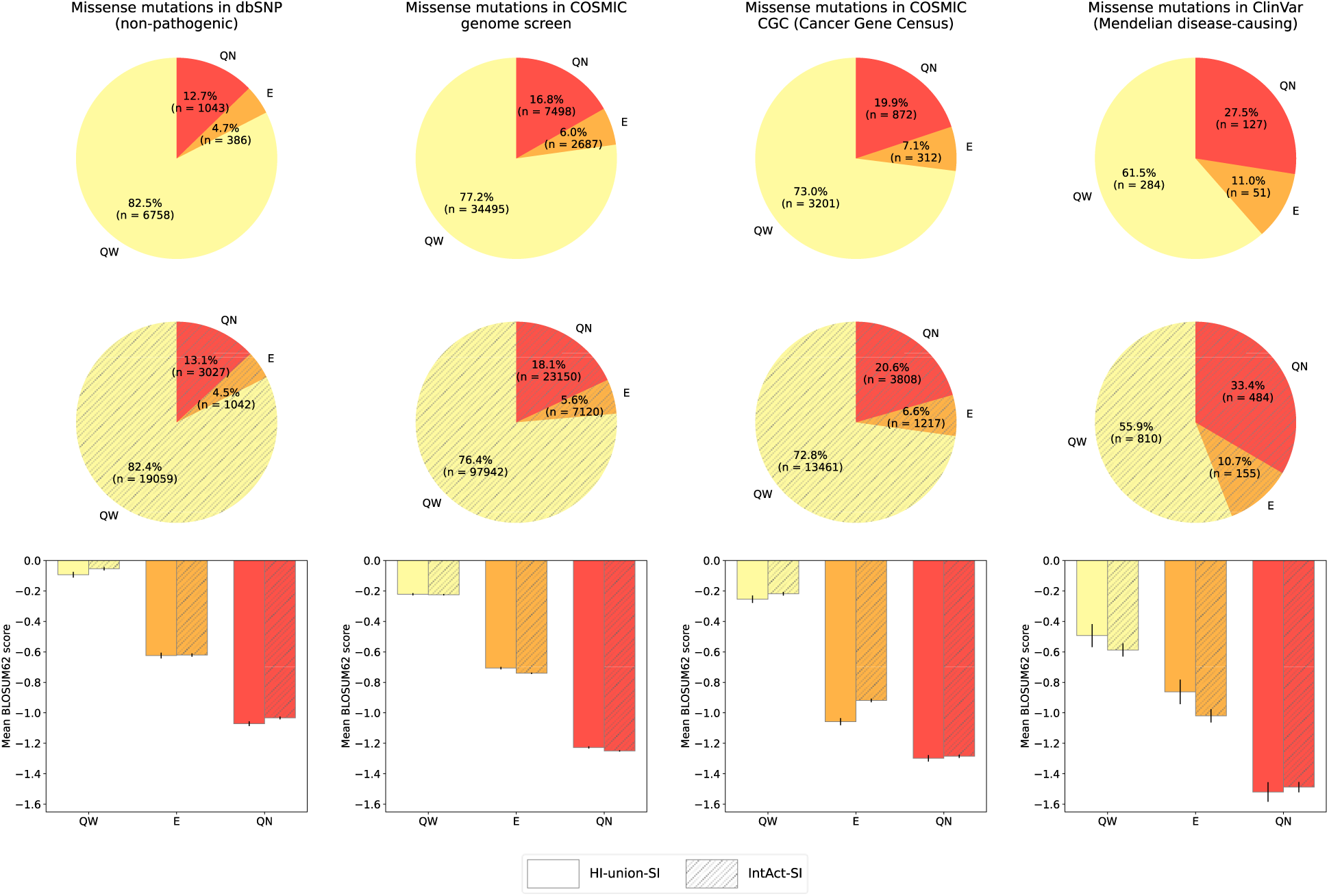
Edgotyping dbSNP (non-pathogenic), COSMIC genome screen, COSMIC CGC (Cancer Gene Census), and ClinVar (Mendelian disease-causing) missense mutations. Edgotypes: edgetic (E), quasi-null (QN), and quasi-wildtype (QW). Edgotyping results for missense mutations mapped onto the HI-union-SI (top row, unhatched piecharts) and IntAct-SI (middle row, hatched piecharts). Mean BLOSUM62 scores of the amino acid substitutions of E, QN, and QW mutations (bottom row, barplots). Error bars represent the standard error.

### Biophysical destabilization is one key contributor to cancer-driving potential

We used the functional effects of destabilizing mutations in Mendelian (monogenic) diseases as a reference to assess the effects of destabilizing mutations in cancer (polygenic). Within the HI-union-SI, edgetic and quasi-null mutations account for 4.7% and 12.7% in dbSNP, 6.0% and 16.8% in the COSMIC genome screen, 7.1% and 19.9% in the COSMIC CGC, and 11.0% and 27.5% in ClinVar (**Figure 2**, top row). Within the IntAct-SI, edgetic and quasi-null mutations account for 4.5% and 13.1% in dbSNP, 5.6% and 18.1% in the COSMIC genome screen, 6.6% and 20.6% in the COSMIC CGC, and 10.7% and 33.4% in ClinVar (**Figure 2**, middle row). In both SIs, the fractions of edgetic and quasi-null mutations increased progressively across dbSNP, the COSMIC genome screen, and the COSMIC CGC. This trend suggests that missense mutations in the COSMIC genome screen are more likely to be passengers than those in the COSMIC CGC, consistent with the expectation that most genome-wide mutations are neutral and resemble dbSNP non-pathogenic variants. Thus, mutations in the COSMIC CGC, a high-quality curated list of genes with causal evidence in cancer (i.e., cancer-driving genes), have a larger likelihood of conferring cancer-driving potential than those across the entire cancer genome. Furthermore, we observed that approximately one-fourth of mutations in each COSMIC dataset and one-third to one-half of mutations in the ClinVar Mendelian disease-causing dataset are either edgetic or quasi-null (**Figure 2**, top and middle rows). This suggests that a large proportion of cancer-associated mutations are quasi-wildtype mutations that likely lead to gain-of-function (GoF) effects within the interactome, aligning with the notion that cancer is a proliferative disease with network-level LoF (e.g., protein-binding destabilization and protein-folding destabilization) being one of the many factors involved.

To further evaluate the contribution of biophysical destabilization to cancer-driving potential, we explored the differences in the folding ΔΔG distribution of quasi-null mutations and in the binding ΔΔG distribution of edgetic mutations across the four mutation datasets. In both SIs, the mean and median folding ΔΔGs of quasi-null mutations increased progressively across dbSNP, the COSMIC genome screen, the COSMIC CGC, and ClinVar (**Figure 3**, top row), indicating that protein-folding destabilization becomes more pronounced with higher cancer-driving potential. This observation further suggests that Mendelian diseases generally involve stronger protein-folding disruptions than cancer. In both SIs, the mean and median binding ΔΔGs of edgetic mutations increased progressively across dbSNP, the COSMIC genome screen, and the COSMIC CGC (**Figure 3**, bottom row), indicating that protein-binding destabilization becomes more pronounced with higher cancer-driving potential. This trend does not continue from COSMIC to ClinVar edgetic mutations, suggesting that the strength of the PPI disruptions in Mendelian diseases may be comparable to those in cancer; however, given our limited edgetic mutation data, no definitive conclusion can be drawn. Altogether, our results demonstrate that mutations in cancer-driving genes exert stronger structural and functional implications on protein folding and binding than mutations across the entire cancer genome as well as non-pathogenic mutations, suggesting that biophysical destabilization is a key contributor to cancer-driving potential.

**Figure 3.**
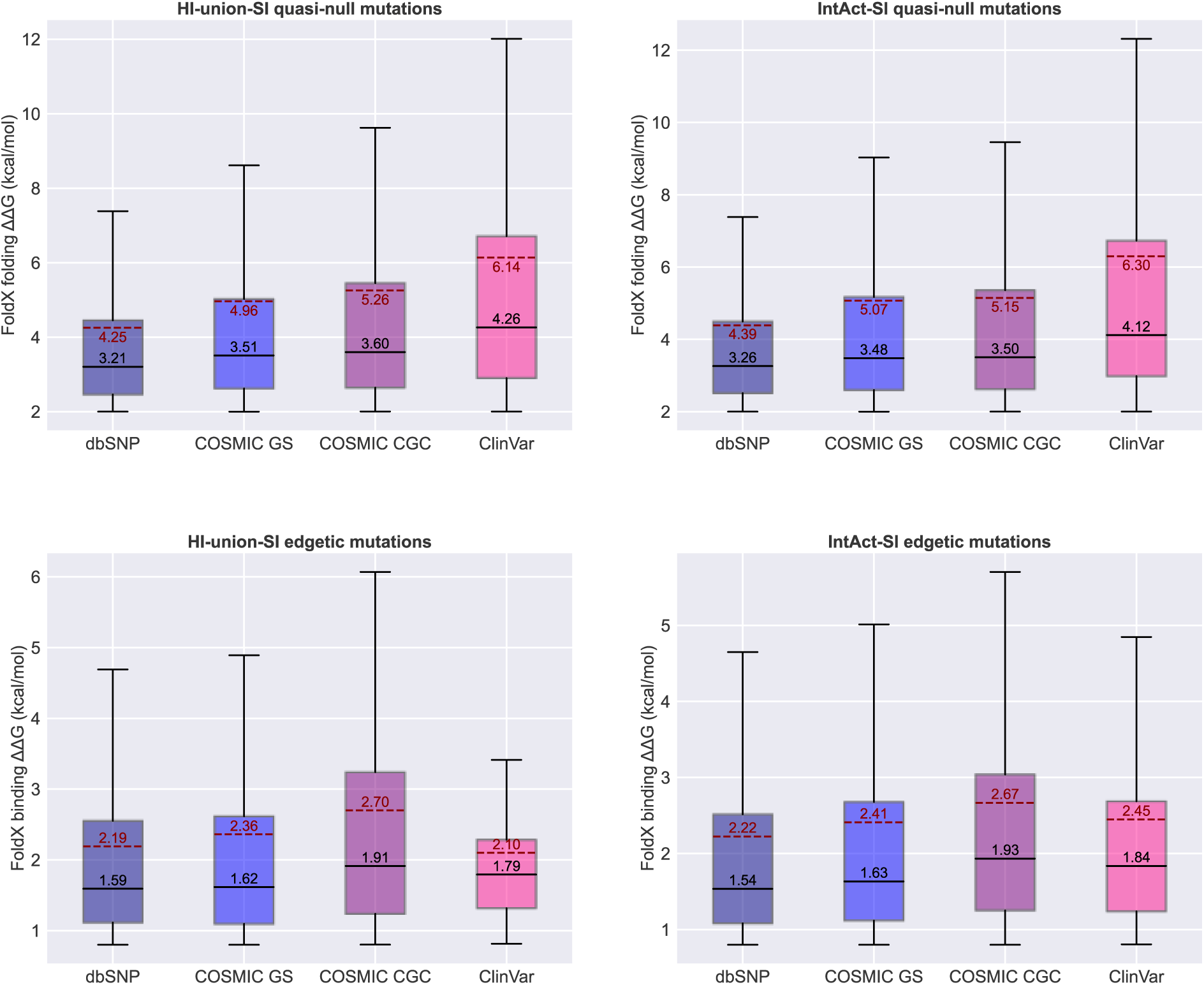
Distribution of folding ΔΔG (protein-folding destabilization) caused by quasi-null mutations and binding ΔΔG (protein-binding destabilization) caused by edgetic mutations. Mutation datasets: dbSNP (non-pathogenic), COSMIC GS (genome screen), COSMIC CGC (Cancer Gene Census), and ClinVar (Mendelian disease-causing). Distributions of ΔΔG for quasi-null (top row) and edgetic (bottom row) mutations mapped onto the HI-union-SI (left column) and IntAct-SI (right column). Mean ΔΔGs are shown by solid black lines and black text, while median ΔΔGs are shown by dotted red lines and red text.

### Quantitative relationship between biophysical destabilization and cancer-driving potential

To evaluate the connection between biophysical destabilization and cancer-driving potential, we calculated the functional and phenotypic deleteriousness of destabilizing mutations using our “fold difference” metric (Su & Xia, 2025). For quasi-null mutations, we investigated the impact of folding ΔΔG on the fold difference between the fractions of quasi-null mutations in datasets of pathogenic (Mendelian disease-causing or cancer-associated) and non-pathogenic missense mutations. Similarly, for edgetic mutations, we investigated the impact of binding ΔΔG on the fold difference between the fractions of edgetic mutations in datasets of pathogenic (Mendelian disease-causing or cancer-associated) and non-pathogenic missense mutations.

In both SIs, the quasi-null and edgetic mutations in ClinVar exhibit larger fold differences across all folding and binding ΔΔG thresholds than those in each of the two COSMIC datasets (**Figure 4**), suggesting that the deleterious effects of biophysical destabilization are milder in cancer than in Mendelian diseases. This observation aligns with the concept that polygenic diseases often require multiple hits to result in pathogenic phenotypes, while monogenic diseases are often associated with one-hit (for dominant traits) or two-hits (for recessive traits) (Karaca et al., 2018; Stratton, Campbell, & Futreal, 2009; Yi et al., 2017). Altogether, these findings indicate that the strong network-level LoF consequences observed in Mendelian diseases can serve as a reference for investigating similar, but generally milder, effects in cancer.

**Figure 4.**
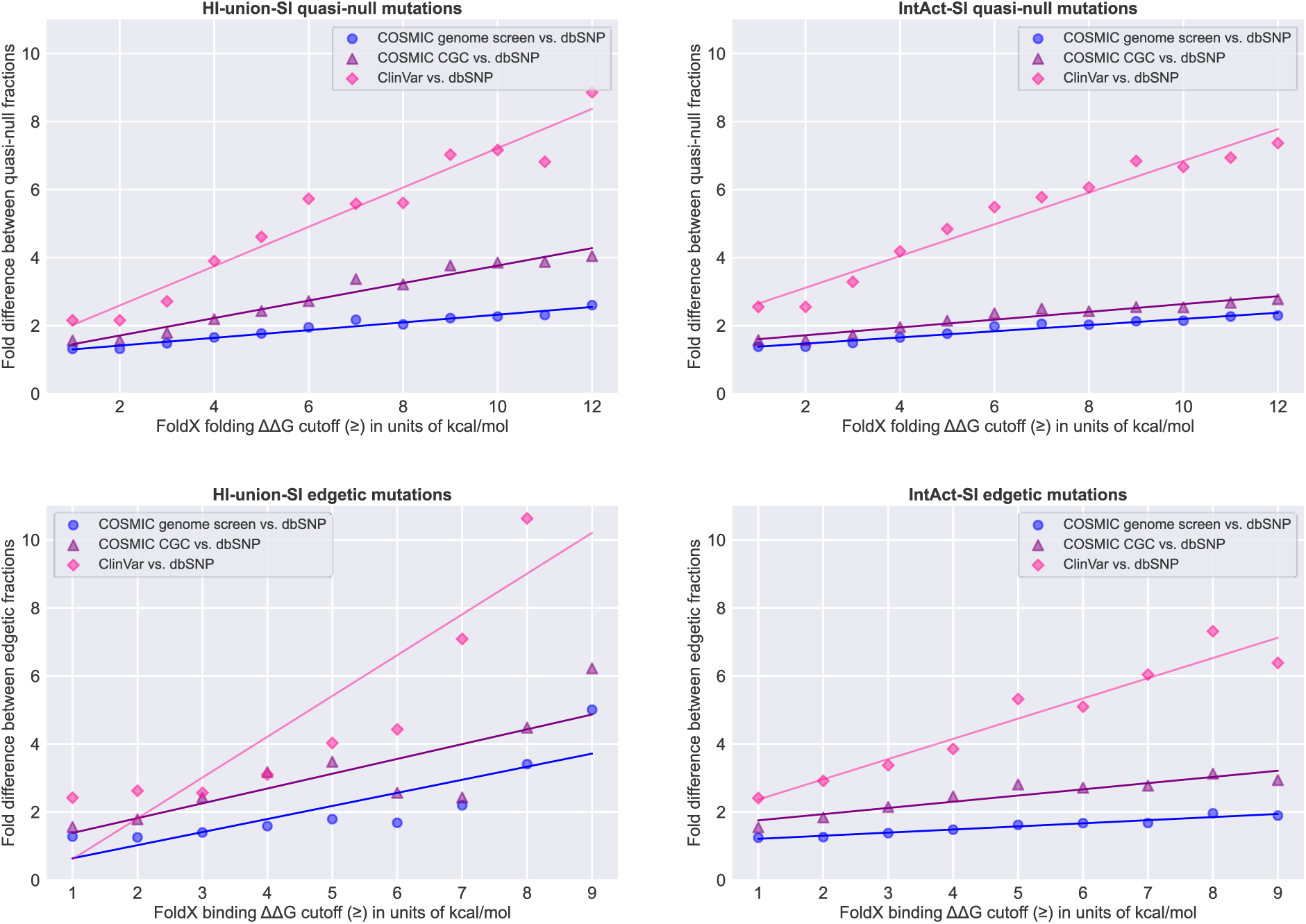
Positive correlation between biophysical destabilization and cancer driving potential (as measured by fold difference). Investigating the cancer-driving potential of quasi-null (top row) and edgetic (bottom row) mutations mapped onto the HI-union-SI (left column) and IntAct-SI (right column). Folding ΔΔG cutoffs for quasi-null mutations ranged from 1 kcal/mol (mildly unstable folding) to 12 kcal/mol (unstable folding). Binding ΔΔG cutoffs for edgetic mutations ranged from 1 kcal/mol (mildly unstable binding) to 9 kcal/mol (unstable binding). Fold differences were calculated using three different pathogenic datasets: COSMIC genome screen (blue dots), COSMIC CGC (Cancer Gene Census, purple triangles), and ClinVar (Mendelian disease-causing, pink diamonds).

For quasi-null mutations in each of the two COSMIC datasets, we observed a positive correlation between the folding ΔΔG threshold (ranging from 1 kcal/mol to 12 kcal/mol) and the fold difference. Specifically in the HI-union-SI, the lines of best fit are y = 0.113x + 1.188 for the COSMIC genome screen and y = 0.257x + 1.195 for the COSMIC CGC (**Figure 4**, top left). In the IntAct-SI, the lines of best fit are y = 0.091x + 1.291 for the COSMIC genome screen and y = 0.115x + 1.488 for the COSMIC CGC (**Figure 4**, top right).

For edgetic mutations in each of the two COSMIC datasets, we observed a positive correlation between the binding ΔΔG threshold (ranging from 1 kcal/mol to 9 kcal/mol) and the fold difference. Specifically in the HI-union-SI, the lines of best fit are y = 0.385x + 0.255 for the COSMIC genome screen and y = 0.435x + 0.949 for the COSMIC CGC (**Figure 4**, bottom left). In the IntAct-SI, the lines of best fit are y = 0.091x + 1.120 for the COSMIC genome screen and y = 0.182x + 1.573 for the COSMIC CGC (**Figure 4**, bottom right).

We further investigated the differences between the COSMIC genome screen and the COSMIC CGC. In both SIs, the quasi-null and edgetic mutations in the COSMIC genome screen consistently exhibit smaller fold differences across all folding and binding ΔΔG thresholds than those in the COSMIC CGC (**Figure 4**), indicating that destabilizing mutations in the CGC are more deleterious and more likely to be cancer-driving than those in the genome screen. This aligns with the notion that most genome-wide somatic mutations are neutral, validating the applicability of our “fold difference” metric for assessing cancer-driving potential. Overall, our findings suggest a positive linear relationship between biophysical destabilization, of both folding and binding, and cancer-driving potential.

### Interactome-wide cancer-associated mutations are predominantly missense rather than nonsense

To better understand the mutational landscape of cancer and the potential for interactome-wide missense and nonsense mutations to be cancer-driving, we assessed the prominence of missense and nonsense mutations among the nonsynonymous mutations in the four mutation datasets. Our datasets of nonsynonymous mutations excluded nonstop mutations as they are scarce and unable to be mapped onto our SIs. We observed that while ClinVar Mendelian disease-causing nonsynonymous mutations are dominated by nonsense mutations (43.0% to 53.8%), both COSMIC cancer-associated (6.1% to 8.2%) and dbSNP non-pathogenic (1.6% to 2.0%) nonsynonymous mutations are unlikely to be nonsense (**Figure 5A**). Furthermore, there is a minor difference in the fractions of nonsense mutations between the COSMIC genome screen (6.1% to 6.3%) and the COSMIC CGC (7.1% to 8.2%) (**Figure 5A**, middle two columns), suggesting that interactome-wide cancer-associated mutations are primarily missense rather than nonsense. Although the proportion of missense and nonsense mutations differs depending on cancer type, our observation is consistent with previous studies showing that missense mutations are the most dominant type of genetic variation across all human cancers (Shi & Moult, 2011; Yi et al., 2017).

**Figure 5.**
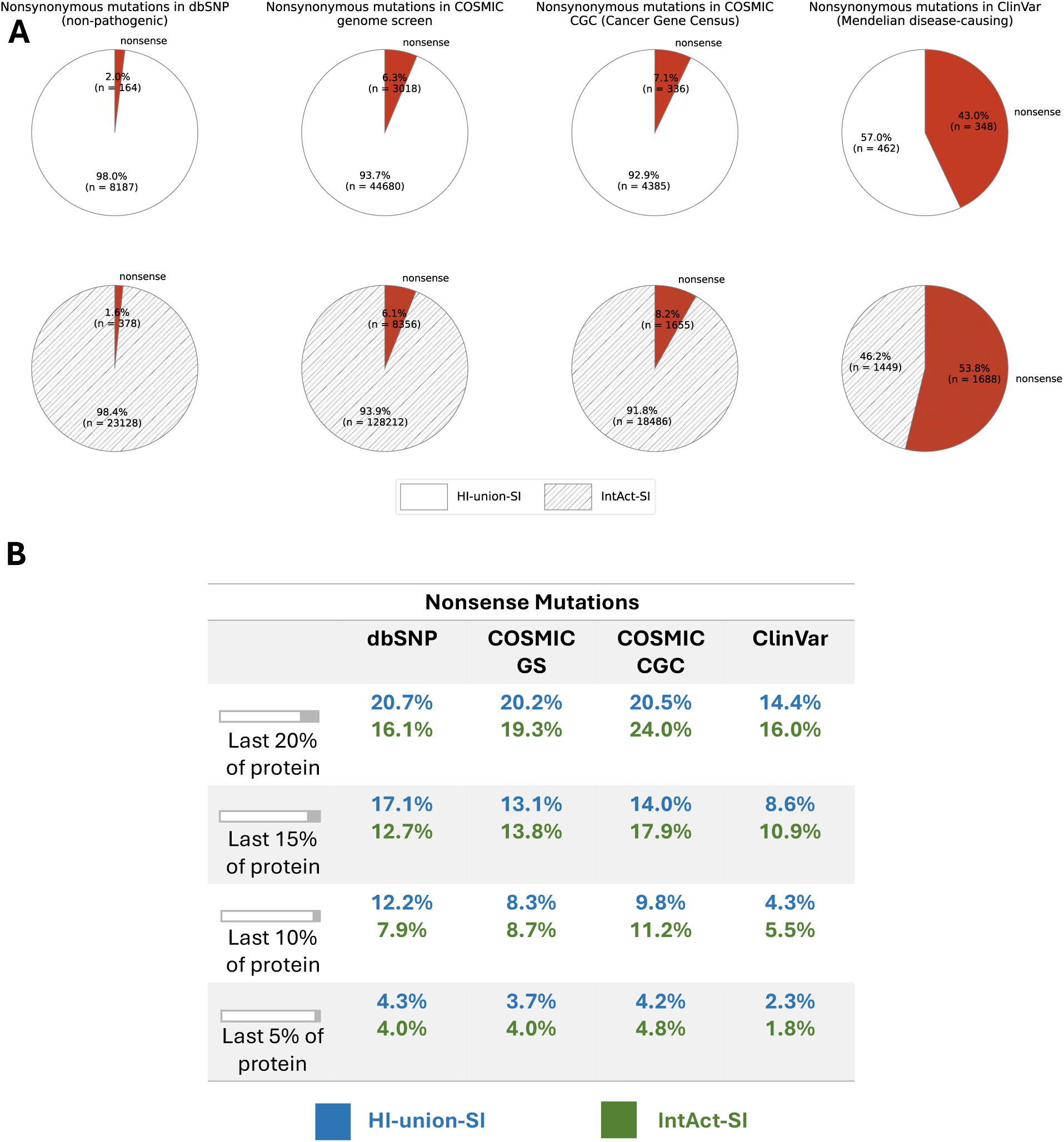
Interactome-wide cancer-associated nonsense mutations are scarce and more likely to be distributed in the tail ends of proteins. Mutation datasets: dbSNP (non-pathogenic), COSMIC GS (genome screen), COSMIC CGC (Cancer Gene Census), and ClinVar (Mendelian disease-causing). **(A)** Proportion of nonsense mutations among the nonsynonymous mutations mapped onto the HI-union-SI (top row) and IntAct-SI (bottom row). Nonsynonymous mutations comprise missense and nonsense mutations; nonstop mutations were excluded due to scarcity and the inability to map them onto proteins. **(B)** Distribution of nonsense mutations in the tail ends of proteins in the HI-union-SI (blue) and IntAct-SI (green). We focused on: the last 20% of proteins, the last 15% of proteins, the last 10% of proteins, and the last 5% of proteins.

### Interactome-wide cancer-associated nonsense mutations are more likely to be distributed in the tail ends of proteins

To further assess the potential for the network-level LoF effects of interactome-wide nonsense mutations to be cancer-driving, we investigated the distributions of COSMIC genome screen and COSMIC CGC mutations in the tail ends of proteins in our SIs. In both SIs, the proportions of COSMIC genome screen and COSMIC CGC nonsense mutations in the last 20%, last 15%, last 10%, and last 5% of proteins were nearly identical (**Figure 5B**, middle two columns). Nonsense mutations in the tail ends of proteins are unlikely to result in complete LoF as most of the proteins are still intact, suggesting that the network-level LoF effects of cancer-associated nonsense mutations may be minimal. Furthermore, the distributions of the COSMIC genome screen and COSMIC CGC nonsense mutations in the tail ends resemble those of the dbSNP non-pathogenic dataset and are approximately 1.5–2 times larger than those of the ClinVar Mendelian disease-causing dataset (**Figure 5B**). This further suggests that the interactome-wide LoF consequences of nonsense mutations may be less severe in cancer than in Mendelian diseases.

## Discussion

Cancer-associated mutations can induce LoF consequences within the human protein interactome by disrupting PPIs and compromising individual protein stability. Here, we structurally resolved the human protein interactome at the atomic level and mapped non-pathogenic, cancer-associated, and Mendelian disease-causing missense and nonsense mutations across the interactome. To assess the biophysical implications of the mutations, we determined their structural locations and performed free-energy calculations. We focused on the network-level LoF effects caused by “edgetic” (binding-destabilizing) and “quasi-null” (folding-destabilizing) missense mutations. We extended our “fold difference” metric, previously proposed for quantifying the functional and phenotypic deleteriousness of LoF consequences in Mendelian diseases (Su & Xia, 2025), to study similar consequences in cancer. We observed a positive correlation between biophysical destabilization and cancer-driving potential. We further observed that destabilizing mutations within cancer-driving genes cause more severe structural and functional consequences than those across the entire cancer genome. This observation aligns with the expectation that cancer-driving genes are likely to harbor more driver mutations than the average gene in the cancer genome, suggesting that driver mutations may be distinguished by their biophysical LoF effects. Our findings suggest that our fold difference metric serves as a valuable quantification for assessing the cancer-driving potential of LoF effects within the interactome. Altogether, our study indicates that biophysical destabilization, of both binding and folding, is a key contributor to cancer-driving potential. Overall, our study of interactome-wide destabilizing mutations provides a mechanistic understanding of the complex processes driving oncogenesis, demonstrating that disruptions to protein folding and binding stability can propagate through the interactome and contribute to cancer-associated phenotypes.

Although interpreting the functional roles of proteins harboring destabilizing mutations would provide a more comprehensive analysis, we were limited by incomplete annotations of tumor suppressor genes (TSGs) and oncogenes. Based on a previous study showing that missense mutations in TSGs primarily destabilize protein folding, whereas those in oncogenes act through alternative mechanisms (e.g., destabilizing less active conformations) (Shi & Moult, 2011), we hypothesize that TSGs are likely to carry a larger proportion of quasi-null mutations than oncogenes. As for edgetic mutations, the mechanisms behind PPI destabilization are more diverse and it remains unclear whether TSGs or oncogenes would harbor a larger fraction of them. The expansion of functional annotations for cancer-associated genes would enable future investigations of the differences in quasi-null and edgetic mutations between TSGs and oncogenes.

Apart from LoF effects, cancer-associated mutations can induce GoF and dominant-negative (DN) effects within the protein interactome. DN effects occur when a mutated protein interferes with the function of the wildtype protein in the same cell, often through the formation of nonfunctional complexes or by binding to and inactivating shared interaction partners (Bergendahl et al., 2019; Veitia, 2009). Mutations associated with GoF and DN effects have different structural and functional consequences from those associated with LoF effects (Gerasimavicius, Livesey, & Marsh, 2022). In particular, the missense mutations we predicted to be “quasi-wildtype” could potentially give rise to GoF or DN effects. Future studies investigating the GoF and DN consequences of interactome-wide mutations would greatly enhance our understanding of the cancer-driving potential of alternative molecular mechanisms.

## Conclusions

In conclusion, our study provides a biophysical perspective on the structural and functional effects of interactome-wide destabilizing mutations and their potential to contribute to oncogenesis. We structurally resolved the human protein interactome at the atomic level and performed a ΔΔG-based assessment of interactome-wide cancer-associated mutations. We applied a simple and universal “fold difference” metric for estimating the cancer-driving potential of folding-destabilizing (“quasi-null”) and binding-destabilizing (“edgetic”) missense mutations. Our observation of the strong positive correlation between biophysical destabilization and cancer-driving potential suggests that disruptions to protein folding and binding stability can propagate through the interactome and contribute to cancer-associated phenotypes. Overall, our study presents important functional insights into interactome-wide destabilizing mutations, providing a mechanistic understanding of the complex processes driving oncogenesis and enabling potential targeted therapeutic strategies.

## Materials and Methods

### Constructing human structural protein interactomes

We built two human SIs, one based on experimentally determined PPIs (HI-union-SI) and the other based on literature-curated PPIs (IntAct-SI), by integrating experimental 3D structures from the PDB (Berman et al., 2000) and homology models for these known PPIs. We first constructed reference interactomes consisting of binary PPIs between Swiss-Prot reviewed proteins using the HI-union (Luck et al., 2020), IntAct (del Toro et al., 2022), and UniProtKB/Swiss-Prot (The UniProt Consortium, 2023) databases. We verified binary PPIs by checking that interacting pairs are homologous to chains in the same PDB complex and that interfacial residues exist between the interacting PDB chains. We then employed homology modeling to build 3D structural models of PPIs in the reference interactomes using BLASTP (Camacho et al., 2009), MODELLER (Webb & Sali, 2016), and the PDB (Berman et al., 2000), as detailed in the computational pipeline established in our previous work (Su & Xia, 2025). Interfacial residues in the 3D homology models were determined using a Euclidean distance-based approach, where a residue was considered interfacial if at least one of its atoms lay within 5 Å of an atom in any residue of the opposing chain. The 3D PPI structural models of the HI-union and IntAct reference interactomes collectively made up our HI-union-SI and IntAct-SI, respectively.

### Processing non-pathogenic, cancer-associated, and Mendelian disease-causing mutations

We obtained missense and nonsense mutations in the GRCh38 genome assembly from the dbSNP database of molecular polymorphisms (Sherry et al., 2001), the COSMIC database of somatic mutations in human cancers (Sondka et al., 2018, 2024), and the ClinVar database of clinically significant variants (Landrum et al., 2016). From the dbSNP database, we selected non-pathogenic missense and nonsense mutations with available 1000 Genomes allele frequencies (Byrska-Bishop et al., 2022) and excluded those annotated with the following phenotypic assertions: conflicting interpretations of pathogenicity, risk factor, pathogenic, likely pathogenic, and drug response. From the COSMIC database, we selected genome screen (COSMIC genome screen) and Cancer Gene Census (COSMIC CGC) somatic cancer-associated missense and nonsense mutations across all nuclear chromosomes. From the ClinVar database, we selected Mendelian disease-causing germline missense and nonsense mutations that were directly labeled as “pathogenic” with supporting evidence and no conflicting phenotypic interpretations.

### Predicting missense mutations that destabilize protein folding (quasi-null) or binding (edgetic)

We mapped the dbSNP non-pathogenic, COSMIC genome screen, COSMIC CGC, and ClinVar missense and nonsense mutations onto our HI-union-SI and IntAct-SI using mRNA to protein mappings and sequences from the Ensembl (Martin et al., 2023) and RefSeq (O’Leary et al., 2016) databases. Mutations were mapped from Ensembl and RefSeq proteins onto Swiss-Prot proteins using their flanking sequences (the adjacent 10 amino acids on either side of the mutation). Within each SI, we selected one mutation for each unique amino acid substitution at any given position in the Swiss-Prot proteins.

We predicted the edgotype (quasi-wildtype, edgetic, quasi-null) (Sahni et al., 2013) of each missense mutation based on its structural location and the mutation-induced change in binding or folding free energy. Due to its computational efficiency and widespread applicability in mutagenesis studies, we used FoldX version 5.1 (Delgado et al., 2025) to perform our free-energy calculations. We first repaired our 3D PPI structural models using the RepairPDB command with default parameters. If a missense mutation coincided with an interfacial residue of any of the protein’s PPIs, we calculated the impact it has on the binding stability (binding ΔΔG) of those PPIs, using the Pssm command with default parameters. We chose a binding ΔΔG threshold of 0.8 kcal/mol, corresponding to the FoldX estimated ΔΔG error (Delgado et al., 2025; Guerois, Nielsen, & Serrano, 2002). If the missense mutation is then found to disrupt one or more PPIs (binding ΔΔG >0.8 kcal/mol), we predicted the mutation to be “edgetic” (binding-destabilizing). All other mutations were categorized as “non-edgetic” mutations, and we calculated the impact that these mutations have on the folding stability (folding ΔΔG) of the proteins, using the BuildModel command with default parameters. If a non-edgetic mutation disrupts the folding stability of the protein (folding ΔΔG ≥2 kcal/mol), we predicted the mutation to be “quasi-null” (folding-destabilizing). The relative solvent accessibility (RSA), which indicates the degree of burial of the mutation, was not considered as we previously showed that RSA is auxiliary in the definition of a quasi-null mutation (Su & Xia, 2025). Conversely, if a non-edgetic mutation maintains the folding stability of the protein (folding ΔΔG <2 kcal/mol), we predicted the mutation to be “quasi-wildtype” (non-destabilizing).

### Fold difference calculations for quasi-null and edgetic mutations

We calculated fold differences, quantitative estimations of deleteriousness or cancer-driving potential, for quasi-null and edgetic mutations, as proposed in our previous study (Su & Xia, 2025). For quasi-null mutations (folding ΔΔG ≥2 kcal/mol), the fold difference is defined as the ratio between the fractions of quasi-null mutations in datasets of pathogenic (COSMIC genome screen, COSMIC CGC, or ClinVar) and non-pathogenic (dbSNP) missense mutations. Similarly, for edgetic mutations (mutations that disrupt at least one PPI, binding ΔΔG >0.8 kcal/mol), the fold difference is defined as the ratio between the fractions of edgetic mutations in datasets of pathogenic (COSMIC genome screen, COSMIC CGC, or ClinVar) and non-pathogenic (dbSNP) missense mutations.

## Code Availability

The scripts written to conduct this study are publicly available on GitHub at https://github.com/tingyisu/CancerLoF.

## Funding

This work was supported by grants from the Natural Sciences and Engineering Research Council of Canada [RGPIN-2019-05952 to Y.X., RGPAS-2019-00012 to Y.X.]; the Canada Foundation for Innovation [JELF-33732 to Y.X., IF-33122 to Y.X.], and the Canada Research Chairs program [CRC-2022-00424 to Y.X.]. T.-Y.S. is supported by a doctoral fellowship from the Fonds de recherche du Québec – Nature et technologies (FRQNT).

## Notes

### Competing Interest Statement

The authors have declared no competing interest.

## References

1. Bergendahl, L. T., Gerasimavicius, L., Miles, J., Macdonald, L., Wells, J. N., Welburn, J. P. I., & Marsh, J. A. (2019). The role of protein complexes in human genetic disease. Protein Science, 28(8), 1400–1411. 10.1002/pro.3667

2. Berman, H. M., Westbrook, J., Feng, Z., Gilliland, G., Bhat, T. N., Weissig, H., … Bourne, P. E. (2000). The Protein Data Bank. Nucleic Acids Research, 28(1), 235–242. 10.1093/nar/28.1.235

3. Byrska-Bishop, M., Evani, U. S., Zhao, X., Basile, A. O., Abel, H. J., Regier, A. A., … Zody, M. C. (2022). High-coverage whole-genome sequencing of the expanded 1000 Genomes Project cohort including 602 trios. Cell, 185(18), 3426–3440.e19. 10.1016/j.cell.2022.08.004

4. Camacho, C., Coulouris, G., Avagyan, V., Ma, N., Papadopoulos, J., Bealer, K., & Madden, T. L. (2009). BLAST+: Architecture and applications. BMC Bioinformatics, 10, 421. 10.1186/1471-2105-10-421

5. Chillón-Pino, D., Badonyi, M., Semple, C. A., & Marsh, J. A. (2024). Protein structural context of cancer mutations reveals molecular mechanisms and candidate driver genes. Cell Reports, 43(11). 10.1016/j.celrep.2024.114905

6. Delgado, J., Reche, R., Cianferoni, D., Orlando, G., van der Kant, R., Rousseau, F., … Serrano, L. (2025). FoldX Force Field revisited, an improved version. Bioinformatics, btaf064. 10.1093/bioinformatics/btaf064

7. del Toro, N., Shrivastava, A., Ragueneau, E., Meldal, B., Combe, C., Barrera, E., … Hermjakob, H. (2022). The IntAct database: Efficient access to fine-grained molecular interaction data. Nucleic Acids Research, 50(D1), D648–D653. 10.1093/nar/gkab1006

8. Engin, H. B., Kreisberg, J. F., & Carter, H. (2016). Structure-Based Analysis Reveals Cancer Missense Mutations Target Protein Interaction Interfaces. PLoS ONE, 11(4), e0152929. 10.1371/journal.pone.0152929

9. Fu, H., Mo, X., & Ivanov, A. A. (2025). Decoding the functional impact of the cancer genome through protein–protein interactions. Nature Reviews Cancer, 25(3), 189–208. 10.1038/s41568-024-00784-6

10. Gerasimavicius, L., Livesey, B. J., & Marsh, J. A. (2022). Loss-of-function, gain-of-function and dominant-negative mutations have profoundly different effects on protein structure. Nature Communications, 13(1), 3895. 10.1038/s41467-022-31686-6

11. Guerois, R., Nielsen, J. E., & Serrano, L. (2002). Predicting Changes in the Stability of Proteins and Protein Complexes: A Study of More Than 1000 Mutations. Journal of Molecular Biology, 320(2), 369–387. 10.1016/S0022-2836(02)00442-4

12. Haque, B., Cheerie, D., Pan, A., Curtis, M., Nalpathamkalam, T., Nguyen, J., … Costain, G. (2025). Leveraging cancer mutation data to inform the pathogenicity classification of germline missense variants. PLOS Genetics, 21(1), e1011540. 10.1371/journal.pgen.1011540

13. Henikoff, S., & Henikoff, J. G. (1992). Amino acid substitution matrices from protein blocks. Proceedings of the National Academy of Sciences, 89(22), 10915–10919. 10.1073/pnas.89.22.10915

14. Kaminker, J. S., Zhang, Y., Waugh, A., Haverty, P. M., Peters, B., Sebisanovic, D., … Zhang, Z. (2007). Distinguishing Cancer-Associated Missense Mutations from Common Polymorphisms. Cancer Research, 67(2), 465–473. 10.1158/0008-5472.CAN-06-1736

15. Karaca, E., Posey, J. E., Akdemir, Z. C., Pehlivan, D., Harel, T., Jhangiani, S. N., … Lupski, J. R. (2018). Phenotypic Expansion Illuminates Multilocus Pathogenic Variation. Genetics in Medicine : Official Journal of the American College of Medical Genetics, 20(12), 1528–1537.10.1038/gim.2018.33

16. Kumar, S., Warrell, J., Li, S., McGillivray, P. D., Meyerson, W., Salichos, L., … Gerstein, M. B. (2020). Passenger Mutations in More Than 2,500 Cancer Genomes: Overall Molecular Functional Impact and Consequences. Cell, 180(5), 915–927.e16. 10.1016/j.cell.2020.01.032

17. Landrum, M. J., Lee, J. M., Benson, M., Brown, G., Chao, C., Chitipiralla, S., … Maglott, D. R. (2016). ClinVar: Public archive of interpretations of clinically relevant variants. Nucleic Acids Research, 44(D1), D862–D868. 10.1093/nar/gkv1222

18. Luck, K., Kim, D.-K., Lambourne, L., Spirohn, K., Begg, B. E., Bian, W., … Calderwood, M. A. (2020). A reference map of the human binary protein interactome. Nature, 580(7803), 402–408. 10.1038/s41586-020-2188-x

19. Martin, F. J., Amode, M. R., Aneja, A., Austine-Orimoloye, O., Azov, A. G., Barnes, I., … Flicek, P. (2023). Ensembl 2023. Nucleic Acids Research, 51(D1), D933–D941. 10.1093/nar/gkac958

20. Mehnert, M., Ciuffa, R., Frommelt, F., Uliana, F., van Drogen, A., Ruminski, K., … Aebersold, R. (2020). Multi-layered proteomic analyses decode compositional and functional effects of cancer mutations on kinase complexes. Nature Communications, 11(1), 3563. 10.1038/s41467-020-17387-y

21. Melamed, R. D., Emmett, K. J., Madubata, C., Rzhetsky, A., & Rabadan, R. (2015). Genetic similarity between cancers and comorbid Mendelian diseases identifies candidate driver genes. Nature Communications, 6(1), 7033. 10.1038/ncomms8033

22. Nishi, H., Tyagi, M., Teng, S., Shoemaker, B. A., Hashimoto, K., Alexov, E., … Panchenko, A. R. (2013). Cancer Missense Mutations Alter Binding Properties of Proteins and Their Interaction Networks. PLOS ONE, 8(6), e66273. 10.1371/journal.pone.0066273

23. O’Leary, N. A., Wright, M. W., Brister, J. R., Ciufo, S., Haddad, D., McVeigh, R., … Pruitt, K. D. (2016). Reference sequence (RefSeq) database at NCBI: Current status, taxonomic expansion, and functional annotation. Nucleic Acids Research, 44(D1), D733–D745. 10.1093/nar/gkv1189

24. Ostroverkhova, D., Przytycka, T. M., & Panchenko, A. R. (2023). Cancer driver mutations: Predictions and reality. Trends in Molecular Medicine, 29(7), 554–566. 10.1016/j.molmed.2023.03.007

25. Porta-Pardo, E., Valencia, A., & Godzik, A. (2020). Understanding oncogenicity of cancer driver genes and mutations in the cancer genomics era. FEBS Letters, 594(24), 4233–4246. 10.1002/1873-3468.13781

26. Rahman, N. (2014). Realising the Promise of Cancer Predisposition Genes. Nature, 505(7483), 302–308. 10.1038/nature12981

27. Sahni, N., Yi, S., Zhong, Q., Jailkhani, N., Charloteaux, B., Cusick, M. E., & Vidal, M. (2013). Edgotype: The link between genotype and phenotype. Current Opinion in Genetics & Development, 23(6), 649–657. 10.1016/j.gde.2013.11.002

28. Schymkowitz, J., Borg, J., Stricher, F., Nys, R., Rousseau, F., & Serrano, L. (2005). The FoldX web server: An online force field. Nucleic Acids Research, 33(suppl_2), W382–W388. 10.1093/nar/gki387

29. Sharifi Tabar, M., Francis, H., Yeo, D., Bailey, C. G., & Rasko, J. E. J. (2022). Mapping oncogenic protein interactions for precision medicine. International Journal of Cancer, 151(1), 7–19. 10.1002/ijc.33954

30. Sherry, S. T., Ward, M.-H., Kholodov, M., Baker, J., Phan, L., Smigielski, E. M., & Sirotkin, K. (2001). dbSNP: The NCBI database of genetic variation. Nucleic Acids Research, 29(1), 308–311. 10.1093/nar/29.1.308

31. Shi, Z., & Moult, J. (2011). Structural and Functional Impact of Cancer-Related Missense Somatic Mutations. Journal of Molecular Biology, 413(2), 495–512. 10.1016/j.jmb.2011.06.046

32. Sondka, Z., Bamford, S., Cole, C. G., Ward, S. A., Dunham, I., & Forbes, S. A. (2018). The COSMIC Cancer Gene Census: Describing genetic dysfunction across all human cancers. Nature Reviews Cancer, 18(11), 696–705. 10.1038/s41568-018-0060-1

33. Sondka, Z., Dhir, N. B., Carvalho-Silva, D., Jupe, S., Madhumita, McLaren K., … Teague, J. (2024). COSMIC: A curated database of somatic variants and clinical data for cancer. Nucleic Acids Research, 52(D1), D1210–D1217. 10.1093/nar/gkad986

34. Song, S., Koh, Y., Kim, S., Lee, S. M., Kim, H. U., Ko, J. M., … Park, S. (2023). Systematic analysis of Mendelian disease-associated gene variants reveals new classes of cancer-predisposing genes. Genome Medicine, 15(1), 107. 10.1186/s13073-023-01252-w

35. Stratton, M. R., Campbell, P. J., & Futreal, P. A. (2009). The cancer genome. Nature, 458(7239), 719–724. 10.1038/nature07943

36. Su, T.-Y., & Xia, Y. (2025). A quantitative comparison of the deleteriousness of missense and nonsense mutations using the structurally resolved human protein interactome. Protein Science, 34(6), e70155. 10.1002/pro.70155

37. Taeubner, J., Wieczorek, D., Yasin, L., Brozou, T., Borkhardt, A., & Kuhlen, M. (2018). Penetrance and Expressivity in Inherited Cancer Predisposing Syndromes. Trends in Cancer, 4(11), 718–728. 10.1016/j.trecan.2018.09.002

38. The UniProt Consortium. (2023). UniProt: The Universal Protein Knowledgebase in 2023. Nucleic Acids Research, 51(D1), D523–D531. 10.1093/nar/gkac1052

39. Veitia, R. A. (2009). Dominant negative factors in health and disease. The Journal of Pathology, 218(4), 409–418. 10.1002/path.2583

40. Webb, B., & Sali, A. (2016). Comparative Protein Structure Modeling Using MODELLER. Current Protocols in Bioinformatics / Editoral Board, Andreas D. Baxevanis … [et Al.], 54, 5.6.1-5.6.37. 10.1002/cpbi.3

41. Xiong, D., Lee, D., Li, L., Zhao, Q., & Yu, H. (2022). Implications of disease-related mutations at protein–protein interfaces. Current Opinion in Structural Biology, 72, 219–225. 10.1016/j.sbi.2021.11.012

42. Yi, S., Lin, S., Li, Y., Zhao, W., Mills, G. B., & Sahni, N. (2017). Functional variomics and network perturbation: Connecting genotype to phenotype in cancer. Nature Reviews Genetics, 18(7), 395–410. 10.1038/nrg.2017.8

43. Zhang, J., Pei, J., Durham, J., Bos, T., & Cong, Q. (2022). Computed cancer interactome explains the effects of somatic mutations in cancers. Protein Science, 31(12), e4479. 10.1002/pro.4479

44. Zhang, Y., Leung, A. K., Kang, J. J., Sun, Y., Wu, G., Li, L., … Yu, H. (2025). A multiscale functional map of somatic mutations in cancer integrating protein structure and network topology. Nature Communications, 16(1), 975. 10.1038/s41467-024-54176-3

